# Hippocampal Neuropeptide Y_2_ receptor blockade improves spatial memory retrieval and modulates limbic brain metabolism

**DOI:** 10.1101/2021.11.17.469003

**Authors:** Marta Méndez-Couz, Héctor González-Pardo, Jorge L. Arias, Nélida M. Conejo

## Abstract

**Introduction:** The neuropeptide Y (NPY) is broadly distributed in the central nervous system (CNS), and it has been related to neuroprotective functions. NPY seems to be an important component to counteract brain damage and cognitive impairment mediated by drugs of abuse and neurodegenerative diseases, and both NPY and its Y_2_ receptor (Y_2_R) are highly expressed in the hippocampus, critical for learning and memory. We have recently demonstrated its influence on cognitive functions; however, the specific mechanism and involved brain regions where NPY modulates spatial memory by acting on Y_2_R remain unclear.

**Methods:** Here, we examined the involvement of the hippocampal NPY Y_2_R in spatial memory and associated changes in brain metabolism by bilateral administration of the selective antagonist BIIE0246 into the rat dorsal hippocampus. To further evaluate the relationship between memory functions and neuronal activity, we analysed the regional expression of the mitochondrial enzyme cytochrome c oxidase (CCO) as an index of oxidative metabolic capacity in limbic and non-limbic brain regions.

**Results:** The acute blockade of NPY Y_2_R significantly improved spatial memory recall in rats trained in the Morris water maze that matched metabolic activity changes in spatial memory processing regions. Specifically, CCO activity changes were found in the dentate gyrus of the dorsal hippocampus and CA1 subfield of the ventral hippocampus, the infralimbic region of the PFC and the mammillary bodies.

**Conclusions:** These findings suggest that the NPY hippocampal system, through its Y_2_R receptor, influences spatial memory recall (retrieval) and exerts control over patterns of brain activation that are relevant for associative learning, probably mediated by Y_2_R modulation of long-term potentiation and long-term depression.

**Highlights:**

- Under hippocampal Y_2_R antagonism, place preference memory retrieval is enhanced
- Spatial retrieval enhancement under Y_2_R blockade is correlated with changes in regional brain energy metabolism
- Enhanced retrieval associated CCO activity increases in the dorsal DG, while decreasing in the ventral CA1, IL cortex and mammillary bodies
- Y_2_R exert control over patterns of brain activation that are relevant for spatial memory expression

## 1. INTRODUCTION

Neuropeptide Y (NPY) was isolated and characterized three decades ago (Tatemoto, 1982). This neuropeptide is widely distributed in the central nervous system (CNS), and it has been associated with functions as food intake (Balasubramanian et al., 2021; Comeras et al., 2019; Stanley & Leibowitz, 1984), cognitive processes (Bertocchi et al., 2021; Thorsell et al., 2000), neuroprotection (Malva et al., 2012; Silva et al., 2005), drug addiction (Gonçalves et al., 2016) and regulation of stress, anxiety, and resilience [For a review, see Reichmann and Holzer (2016)]. However, the role of NPY as a memory modulator is still controversial (Gøtzsche & Woldbye, 2016). Previous research showed that the NPYergic system participates in cognitive processes either improving or impairing short- and long-term memory, and these effects are claimed to be both dose- and brain region-specific (Bertocchi et al., 2021; Flood et al., 1987; Gøtzsche & Woldbye, 2016; Kornhuber & Zoicas, 2017, 2020; Thomas & Ahlers, 1991).

The CNS NPYergic system exerts its effects through Y_1-_Y_6_ receptors (Parker & Herzog, 1999; Xapelli et al., 2006). All of them are postsynaptic receptors, except Y_2_R, a presynaptic autoreceptor that inhibits NPY release (Decressac & Barker, 2012; Stani et al., 2006; Stanić et al., 2011). The hippocampal Y_2_R, which presynaptically inhibits glutamate release, also undergoes *de novo* expression in mossy fibres and granule cells (Furtinger et al., 2001; Gobbi et al., 1998).

NPY and its receptors are highly expressed in brain regions that contribute to cognitive processing (Parker & Herzog, 1998, 1999). Specifically, Y_2_R is highly expressed in the amygdala, hippocampus and hypothalamus, the brain areas considered essential for contextual memory processes, and their integration with emotional components (Stanić et al., 2011). In the hippocampus, Y_2_R is found at higher levels in the pyramidal cell layers of CA1 and CA3, although is also present in the DG-hilus region(Parker & Herzog, 1999).

Regarding the specific effects of the NPYergic system on learning and memory, a growing body of literature suggests its involvement in the encoding and recall learning phases (Bertocchi et al., 2021; dos Santos et al., 2013; Hörmer et al., 2018; Méndez-Couz et al., 2021). Interestingly, NPY showed a neuroprotective role against methamphetamine-induced excitotoxicity in the hippocampus (Gonçalves et al., 2012) or against oxidative stress, via Y_2_R, preventing depressive behaviour and spatial memory deficits in mice (dos Santos et al., 2013).

For cognitive processing to occur, neurons carry on multiple energy-consuming activities (Wong-Riley, 1989, 2012). Their supply of energy, in form of ATP, comes almost entirely from oxidative phosphorylation, in contrast to astrocytes that rely on glycolysis (Almeida et al., 2004), which make them heavily dependent on mitochondrial function. The brain metabolic state was analysed in this study using a cytochrome c oxidase (CCO) activity analysis. CCO is the last member of the electron transport chain located in the inner mitochondrial membrane [see review by Chicherin et al. (2019)]. The tight coupling between neuronal activity and oxidative energy metabolism converts the CCO activity into a good and useful endogenous metabolic biomarker for neuronal activity (Gonzalez-Lima & Cada, 1994; Gonzalez-Lima & Jones, 1994; Wong-Riley, 1989, 2012). CCO activity can, therefore, be interpreted as a history of cumulative energy demands of brain cells underlying prolonged stimulation or training on behavioural tasks – days or weeks – (Agin et al., 2001; Bertoni-Freddari et al., 2001; Conejo et al., 2010; Hescham et al., 2014); this represents a complementary technique to the more short-lived immediate early genes (IEG) studies (minutes or several hours after evoked neuronal stimulation). CCO quantitative histochemistry has also proved to be a reliable method to measure changes in brain metabolism associated with endocrine alterations and neurodegeneration (Morán et al., 2013) as well as learning and memory processing (Conejo et al., 2010; Gasalla et al., 2016; Méndez-Couz et al., 2016) and, specifically, to measure brain regional changes associated with the retrieval of spatial learning (Conejo et al., 2013; Méndez-Couz et al., 2015a; Zorzo et al., 2021a; 2021b).

Following the stimulation of N-Methyl-D-aspartate (NMDA) receptors, neurons transiently synthesize nitric oxide (NO) in a calcium/calmodulin-dependent manner. NPY modulates the production of NO in the microglia induced by interleukin-1β (IL-1β). As shown by other authors (Ferreira et al., 2010; Malva et al., 2012), NPY treatment inhibits NO production as well as the expression of inducible nitric oxide synthase. Nitric oxide acts as a cellular messenger, activating soluble guanylyl cyclase and participating in the transduction signallin g pathways involving cyclic GMP [See (Moncada & Bolaños, 2006)]. NO production can also act as a pathophysiological mediator, related to heart failure among other diseases (Almeida et al., 2004; Moncada & Bolaños, 2006). In the CNS, high amounts of NO inhibit the mitochondrial CCO in neurons, resulting in neuronal depolarization and calcium-dependent vesicular glutamate release, followed by excitotoxicity. Ultimately, neurons die by excitotoxicity via NMDA (Brown, 2007). NO also binds to CCO, inhibiting cell respiration in a reversible process, competing with oxygen (Moncada & Bolanos, 2006) and leading to the release of superoxide anion from the respiratory chain. Besides, NPY is a known regulator of feeding and energy metabolism by affecting mitochondria that, in turn, act as energy sensors in AgRP/NPY hypothalamic neurons. This stimulates appetite and increases energy oxidative metabolism (Billington & Levine, 1992; Haigh et al., 2020; Su et al., 2016). Therefore, NPY can ultimately modulate the activity of key mitochondrial enzymes involved in cellular respiration and oxidative metabolism like CCO, measured here.

In a previous NPY receptor expression study, we showed changes in spatial memory task performed under serial Y_2_R blockade (Méndez-Couz et al., 2021), associated with NPY Y_1_R and Y_2_R brain expression changes in the dorsal hippocampus and the prefrontal cortex, suggesting a rapid metaplastic regulation of NPY receptors contributing to spatial memory. The specific role of NPY via Y_2_R on spatial orientation memory, including the region-specific involvement and its mechanism of action in this process, remains – nevertheless – unknown.

In the current study, we evaluated the brain regional metabolic activity changes following a spatial memory recall task under acute infusion of a Y_2_R intrahippocampal antagonist, which improved spatial memory associated with the increased metabolic activity of the DG, and decreased metabolic activity in the CA1 subfield of the ventral hippocampus, the infralimbic cortex and mammillary bodies.

## 2. MATERIAL AND METHODS

### 2.1. Animals

Male adult Wistar rats (*Rattus norvegicus)*, weighing between 250–330g, were used (N=30). They were obtained from the University of Seville central animal facilities (Seville, Spain) and housed in a temperature-controlled room (23±2°C). Lighting was kept on a 12-h light/dark cycle with lights on from 08:00–20:00 h. Rats were kept in standard laboratory cages (20 × 35 × 55 cm), with four rats in each cage with *ad libitum* access to food and tap water.

### 2.2 Behavioural procedure

The rats were first tested in a neurological assessment battery to discard possible motor and sensory deficits before and after the surgery. The neurological tests used allowed us to evaluate the following reflexes: abduction response of hind limbs, grasping reflex, extension and flexion reflexes, hearing and vestibular responses, head shaking reflex, pupillary reflex, negative geotactic responses and righting reflex (Bures et al., 1976). Rats were handled daily, 5 days before the surgery, to reduce anxiety-like behaviour related to contact with the experimenters. See the timeline of the experiment in Figure 1.

**Figure 1:**
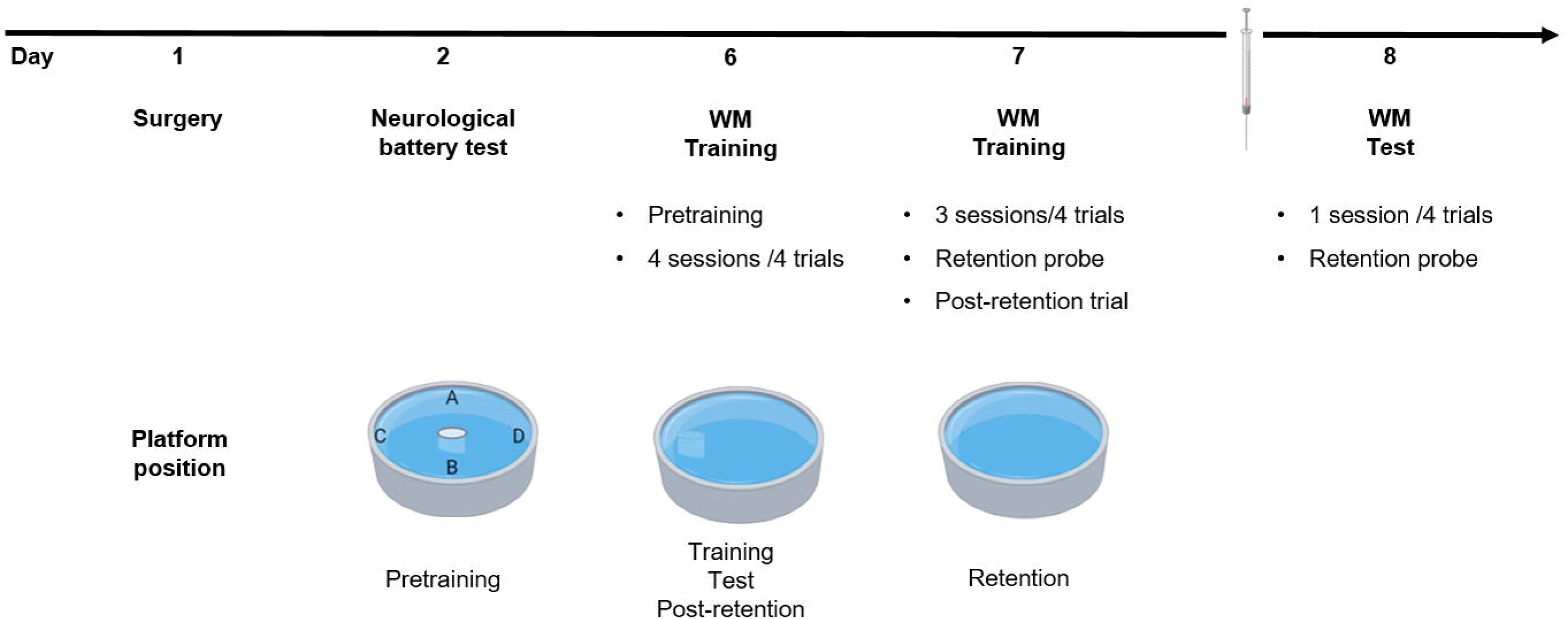
Experiment timeline and behavioural procedure. **Top)** Temporal line of the procedure. **Middle)** After recovery from surgery, rats were trained in a reference spatial memory task in the water maze for two days. On the first day, they went through four sessions of four trials. Meanwhile, on the second day, three sessions of four trials each were carried out. After the last trial of the acquisition, a transfer test, to determine the virtual quadrant preference of the rats, was performed. An additional trial was carried out in which the platform was placed in its original position, to avoid early extinction of the acquired task. One day after the acquisition was complete, rats received intrahippocampal infusion with the Y_2_R selective antagonist (1 nmoL/µL BIIE0246 in 0.9% physiological saline +1% DMSO) or vehicle saline solution, 30 min before the retrieval test in the water maze. Treated rats underwent four trials with the platform in the maze and an additional transfer test without the platform. **Bottom)** Position of the platform in the different stages of the water maze task. In the pretraining phase, the platform was visible, 2 cm above the water surface. During acquisition, the platform was hidden in the maze; rats orientated themselves through distal cues situated in panels situated around the maze. For the retention probe after the acquisition, the platform was removed from the maze. The time to reach the platform (escape latencies) and the time that rats spent in the virtual reinforced quadrant area during the transfer memory probe test were taken as variables related to spatial learning and memory performance.

### 2.3. Surgery

Rodents were deeply anaesthetized with xylazine (5 mg/kg, i.m.) and ketamine (80-100 mg/kg, i.p.), and set in a David Kopf (Tujunga, CA) or Narishige (Japan) stereotaxic frame. Stainless steel cannulae of 22G inner diameter (Becton Dickinson S.A., Spain) were stereotactically implanted bilaterally in the CA1 region of the dorsal hippocampus (coordinates from bregma: AP −3.6, L ±2.6, DV −2.1 mm). Cannulae were fixed to the skull using dental acrylic cement (Glaslonomer Cement, Shofu Inc., UK) and anchor screws. Rats were allowed to recover after surgery for 5 days.

After this resting period, the rats underwent the same neurological assessment battery explained above to discard any possible abnormality caused by the stereotaxic procedure.

### 2.4. Groups

Rats were randomly assigned to one of the three experimental groups as follows:

1. Control cage (CC): rats maintained in their home cage without further pharmacological, surgical or behavioural interventions. This group was added to ensure that brain metabolic activity changes were due to behavioural and pharmacological modifications.
2. Experimental group (Exp): received 1 nmoL/µL BIIE0246 in 0.9% physiological saline containing 1% DMSO (1 µL/hemisphere).
3. Control saline vehicle group (Veh): rats were administered 0.9% physiological saline containing 1% DMSO (1 µL/hemisphere).

### 2.5. Injection

Following the protocol described by Méndez-Couz et al. (2021), rodents received the treatment through the permanent cannula 30 min before the retrieval test was performed on the third day of the Morris water maze (MWM). Rats were randomly assigned to the experimental or vehicle saline group. Solutions were infused at 0.5 µl/min and the microcannula was held inside for an additional 60 s to prevent fluid from backing up into it.

### 2.6. Spatial memory training

#### 2.6.1. Apparatus

The MWM was composed of a circular water tank made of black fibreglass, measuring 1.5 m in diameter by 75 cm in height (Morris, 1984). Following the experimental setting conditions described by Méndez-Couz et al. (2014,2015a), the pool was filled with tap water, and a black escape platform was placed hidden beneath the water surface. The water temperature was kept at 20±1°C during the entire training period. The pool was surrounded by black panels located 40 cm from the maze, on which highly contrasted printed geometric visual cues were placed, acting as allocentric cues. The room was softly illuminated by two halogen spotlights facing the ceiling (500 W). Each trial was recorded and later analysed offline, using a computerized video-tracking system (Ethovision Pro, Noldus Information Technologies, Wageningen, The Netherlands). Measured variables included the time spent to reach the platform (latencies) when the platform was present in the maze and the time spent in each of the four virtual quadrants in which the pool was divided (A, B, C or target, and D) during the transfer tests.

The behavioural protocol in the water maze was applied as described in Méndez-Couz et al. (2021); it is also briefly detailed below (Figure 1).

#### 2.6.2. Pretraining

In order to habituate the rats to the task in the MWM, they received a pretraining session with a visible platform present at the centre of the maze. During this first stage, rats received four trials, in which they were released facing the pool walls following a pseudo-random sequence. The escape platform was set 2 cm above the water surface so that it was visible from afar. Rats could swim up to 60 s to locate the platform in each trial or were gently manually guided to it after that time. Once on the platform, rats were left 15 s on it, and they rested 5 s outside the maze in a plastic container until commencing the next trial.

#### 2.6.3. Spatial reference memory task: training session

During training, the rats were required to locate a hidden platform placed in the centre of a determined quadrant for each rat. Therefore, they had to use external visual cues situated around the pool to perform the task (see Figure 1).

After pretraining, on the first day, rats received four training sessions of four trials each, resting 30 min in their home cage between sessions. In each trial, rats were released from the border of each of the quadrants in a pseudorandom order to search for the hidden escape platform located 1.5 cm beneath the water surface (Figure 1, Bottom). Rats were allowed to swim for 60 s to reach the platform or manually guided to it after this time; once they achieved the platform, they remained there for 15 s. The intertrial time was 5 s, in which they were allowed to rest outside the maze in a plastic container, different from their home cage. On the following day, they received three identical sessions of four trials, until accomplishing on average a learning criterion of 20 s latency to reach the platform; training was stopped at this point to avoid overtraining.

After the last trial on day two, rats were submitted to a retention probe test trial to find out whether they remembered the location of the hidden platform. During this probe, the platform was removed from the pool, and rats were released from the contralateral side to the previously reinforced quadrant. The test lasted for 60 s, after which the rats were placed again in a plastic bucket outside the maze. To prevent the early extinction of the previously learned task, all rats received an additional trial in which the platform was present in the maze in its original place. The latencies during this trial were included for analysis together with the data from the last session of acquisition (Figure 2a).

**Figure 2:**
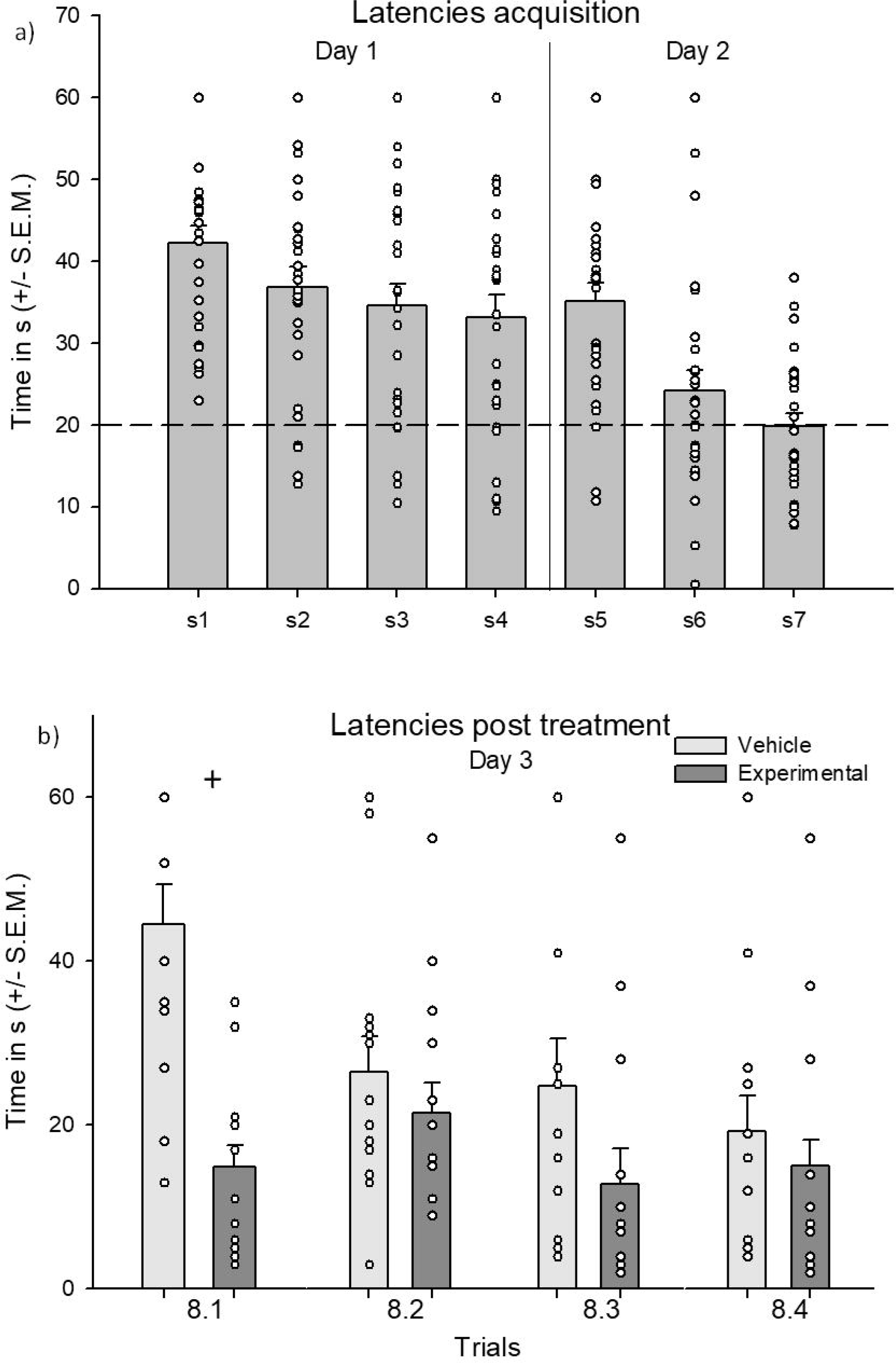
After treatment, rats required less time to reach the platform. **a) Acquisition learning curve.** Rats learned the hidden platform spatial task. Differences along the training sessions 1 to 7 were found, showing a typical learning curve with decreasing mean latencies needed to reach the platform (*p*<0.001). The acquisition was stopped at Session 7 after achieving, on average, the learning criterion of 20 s to reach the platform, represented by a dashed line. **b) Latencies post-treatment.** One day after the task was successfully acquired, the spatial memory test took place, Y_2_R antagonist (for the experimental group) or vehicle saline solution (for the saline control group) was infused 30 min prior to a spatial memory test was performed. Experimental and vehicle groups presented significant differences in the latency needed to reach the platform ^+^(*p*=0.001); specifically, the treated group needed less time to reach the platform as compared to the vehicle saline control in the first trial, although both groups reached low latencies at the end of the retrieval test. The average amount of time spent is represented in bars, dots represent individual data points.

#### 2.6.4. Retrieval test

The day after finishing the second day of acquisition, a retrieval test was performed. Rodents were intracerebrally infused with the drug (1 nmoL/µL BIIE0246 in 0.9% physiological saline + 1% DMSO) or vehicle saline solution 1% DMSO, 30 min before the spatial memory test in the MWM, consisting of four trials in the same conditions as explained above. Additionally, a new memory retention probe test was performed on day three.

### 2.7. Molecular analysis

#### 2.7.1. Infusion site

All animal brains from the experimental and vehicle groups were analysed for the infusion site through serial brain slices section selection and posterior analysis after cresyl violet Nissl staining according to Paxinos and Watson (2004). One animal was discarded due to injection misplacement. The extension of the drug area of influence at the injection site was estimated in previous pilot studies to be around 1.5 mm in diameter, sufficient to cover the targeted CA1 dorsal field of the hippocampus.

#### 2.7.2. Brain cytochrome c oxidase histochemistry

Randomly selected rats from the vehicle and experimental groups, and additional rats selected as CC group (Veh N=6, Exp N=8, CC N=8) were sacrificed 90 min after finishing the behavioural tasks or directly from the cages in the latter case; brains were quickly removed then frozen in isopentane at −70°C (Sigma–Aldrich, Madrid, Spain) and stored at −40°C, to preserve the brain tissue and enzyme activity. Brains were subsequently cut at 30µm-thick coronal sections using a cryostat microtome (Microm International GmbH, model HM 505-E, Heidelberg, Germany). These sections were mounted on gelatinized slides and stored at −40°C until further processing.

A modified version of the quantitative CCO histochemical method developed by Gonzalez-Lima and Jones (1994) was used. Staining variability across different baths was controlled by sets of tissue standards. These standards were obtained from Wistar rat brain homogenates of known CCO activity that were determined spectrophotometrically at different thicknesses (10, 30, 50 and 70 µm). Following the previously described protocol by Conejo et al. (2013), the standards were included with each batch of slides. In short, each set of slides were fixed for 5 min with a 0.5 (v/v) glutaraldehyde solution, rinsed three times in phosphate buffer (PB) and preincubated 5 min in a solution containing 0.05 M Tris buffer pH 7.6 with 275 mg/L cobalt chloride 10% (w/v), sucrose and 0.5 (v/v) dimethylsulfoxide. After the sections had been rinsed in PB (pH 7.6; 0.1 M), they were incubated at 37°C for 1 h in darkness, in a solution containing 50 mg 3,3′-diaminobenzidine, 15 mg cytochrome c (Sigma, St. Louis, MO, USA) and 4 g sucrose per 100 mL PB (pH 7.4; 0.1 M). The reaction was stopped by fixing the tissue in buffered formalin (10% w/v sucrose and 4% formalin) for 30 min at room temperature (RT). After being fixed the slides were dehydrated, cleared with xylene and coverslipped with Entellan (Merck, Darmstadt, Germany).

CCO histochemical staining intensity was measured by densitometric analysis using a computer-assisted image analysis workstation and image analysis software (MCID, InterFocus Imaging Ltd., Linton, England). Twelve measurements of relative optical density were taken per region (4 per slice). To compare and consider possible staining variations across brain sections from different staining baths, measurements were also taken from CCO-stained brain homogenate standards. Regression curves between section thickness and known CCO activity, previously assessed by spectrophotometric assay in each set of standards, were calculated. Lastly, the average relative optical density measured in each brain region was converted into CCO activity units (1 unit: 1 µmol of cytochrome c oxidized/min/g tissue wet weight at 23°C), using the previously calculated regression curve in each homogenate standard. Averages were calculated per region and animal. Regions to analyse included the prelimbic (PL), and infralimbic regions (IL) of the medial prefrontal cortex and primary motor cortex (M1), all of them measured at ^+^/_-_ 3.70 mm from Bregma and the parietal cortex (PAR) ^+^/_-_ −3.80 mm. In addition, the following subcortical regions were also taken: dorsal subfields of hippocampus including CA1, CA3 and dentate gyrus (dCA1, dCA3, dDG) at ^+^/_-_ −3.30 mm and ventral hippocampus subfields (vCA1, vCA3, vDG), taken at ^+^/_-_ −4.52 mm from Bregma. Medial (MeA), basal (BaA), lateral (LaA) and central (CeA) amygdaloid nuclei measured at ^+^/_-_ −3.14 mm from Bregma; medial (MM), lateral (LM) and supramammillary (SuM) nuclei of the mammillary bodies measured at ^+^/_-_ −4.52 mm, as well as the premammillary nucleus (PM) taken at ^+^/_-_ −4.16 mm. The selected brains regions were defined according to Paxinos and Watson’s (2004) atlas.

### 2.8. Statistical analysis

#### 2.8.1. Behavioural tests

A one-way ANOVA test was applied to evaluate the latencies across training sessions during the acquisition days, with “session” as a factor. Two-way repeated-measures ANOVA tests were applied to evaluate the group differences in the escape latencies after the treatment. Afterwards, Holm-Sidak tests were used to further analyse significant interactions between group and training sessions. Holm-Sidak’s post-hoc tests were also used to evaluate differences across training sessions in each experimental group. To analyse the swimming time spent in each virtual quadrant of the maze during the retention probe after the acquisition, a one-way repeated-measures ANOVA was carried out. After the infusion, a two-way RM ANOVA test was used to study these differences in quadrant preference in both experimental groups during the retention probe. Holm-Sidak’s post-hoc tests were used in case of significant ANOVA results.

#### 2.8.2. Cytochrome c oxidase histochemistry measurement

Experimental, vehicle saline and CC groups differences in CCO activity measured in each brain region were analysed by a mixed model one-way ANOVA. Tukey’s post hoc tests were used to assess differences between pairs of experimental groups when ANOVA indicated significant group differences. Significance levels were set to *p*<0.05. Statistical analysis was performed using Sigma-Plot 11 (Systat Software, Chicago, USA).

## 3. RESULTS

### 3.1. Behavioural tests

No rats were discarded due to their neurological reflexes’ responses after the surgical procedure.

#### 3.1.1. Rats acquired the hidden platform spatial memory task

Results showed that rats (N=28) acquired the spatial memory task before treatment, as shown by the significant decrease in the latencies needed to reach the platform through the acquisition sessions one to seven (F_(6,182)_=10.35, *p*<0.001). Specifically, the post-hoc test showed differences between the first and the last acquisition session (Holm-Sidak, *p*<0.001) (Figure 2a). By the end of the second day, rats reached, on average, the learning criterion of 20 s to arrive at the platform, so the acquisition phase was considered finished. Rats that failed to reduce the amount of time needed to reach the platform from the first to the last sessions or presented an inverted learning curve were discarded, and their data were not included in the study (N=2). The acquisition phase was stopped at this point to avoid overtraining, lack of motivation and an early extinction of the previously learned response.

After finishing the acquisition phase, rats underwent a retention probe test, in which a preference for the previously reinforced quadrant was observed (F_3,108_=41.41, *p*<0.001), (Holm-Sidak between reinforced quadrant and rest of the quadrants, *p*<0.001) (Figure 3a).

**Figure 3:**
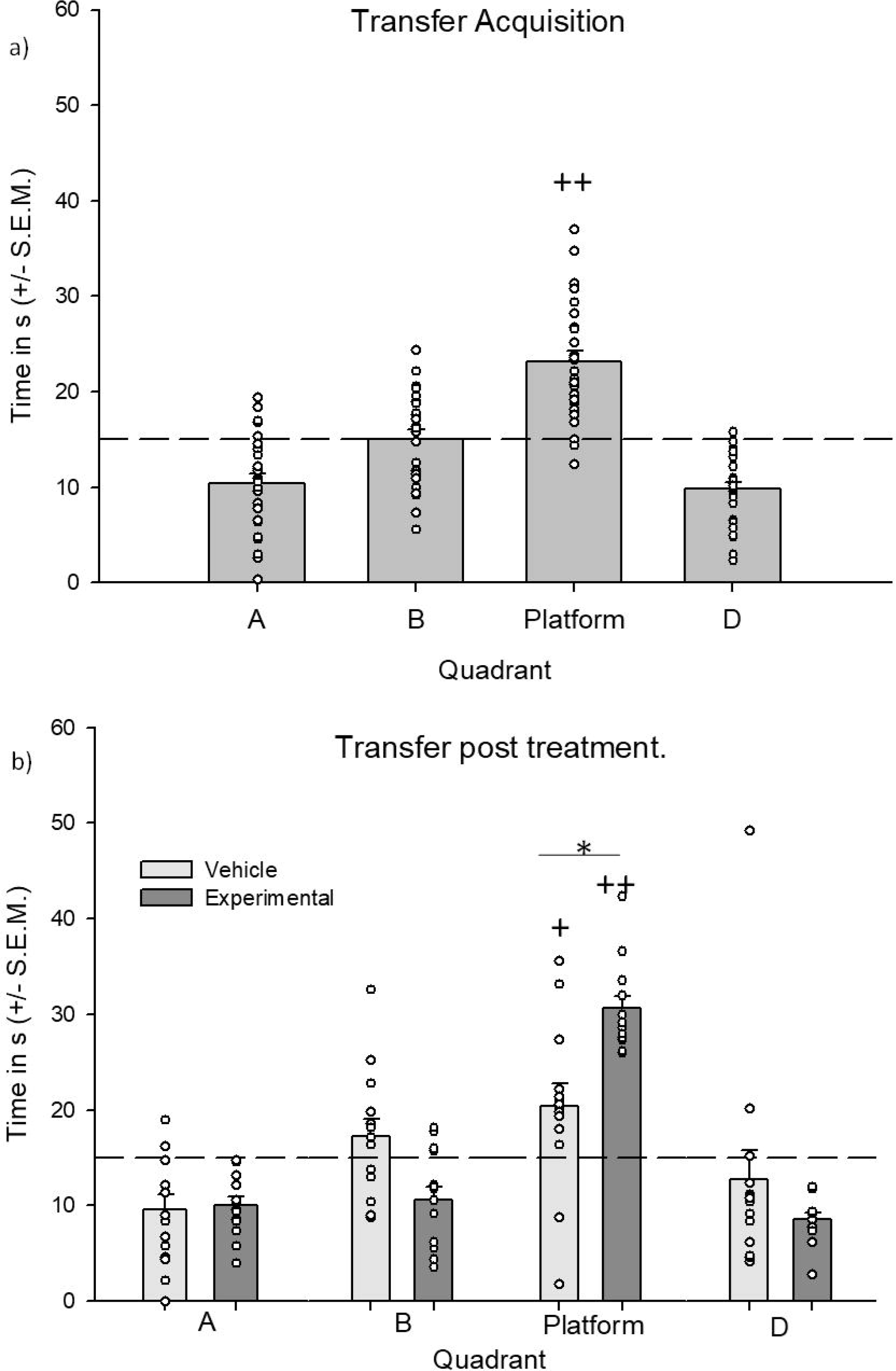
Treated rats spent more time in the reinforced quadrant after treatment than vehicle saline controls. Mean time spent in the virtual quadrants in which the maze was divided **a)** At the end of the acquisition phase, rats spent more time in the previously reinforced quadrant as compared to the rest of the quadrants. **b)** After the saline/drug infusion on the test day, the vehicle saline group spent more time in the reinforced quadrant as compared to A and D; meanwhile, the experimental group spent more time in the reinforced quadrant than in all the rest. When groups were compared, the NPY Y_2_R antagonist treated rats spent more time in the reinforced quadrant as compared to the vehicle control *(*p*<0.05) ++(*p*<0.001) +(*p*<0.01)

#### 3.1.2. Administration of the Y_2_R antagonist improved spatial orientation ability in the MWM test

Rats that had learned the task were randomly assigned to the vehicle or experimental groups. One day after the last acquisition session, they were administered with the saline vehicle or the antagonist solution, respectively, and after 30 min, they were submitted to an early retrieval test consistent in four trials in the MWM. When comparing the latencies to reach the platform through the four post-treatment trials, we found significant differences between vehicle and experimental groups (Two ways RM ANOVA F_(1,76)_=13.58, *p*=0.001). These differences depended on the analysed trial (Two ways RM ANOVA F_(23,76)_=3.37, *p*=0.023). Specifically, the vehicle group was slower to reach the platform as compared to the experimental group during the first trial after the drug administration (Holm-Sidak, *p*=0.001). Eventually, all groups showed improvement, by gradually decreasing latencies to reach the platform over the 4 trials of the test (Trial 8.1 vs Trial 8.4, *p*<0,001). Both groups needed under 20 s on average to reach the platform in the last trial post-treatment (Figure 2a).

The Y_2_R antagonist effect on memory retrieval was also observed in the preference for the reinforced quadrant during the retention probe carried out on the test day. Results show an interaction between group and quadrant factors (F_(3,78)_=4.28; *p=*0.008). Vehicle and experimental groups spent a significantly different amount of time in the different quadrants of the maze on the test day (F_(1,108)_=5.62; *p*=0.025). The analysis also showed differences in the factor quadrant (F_(3,98)_=13.93; *p*<0.001). Specifically, the experimental group preferred the previously reinforced quadrant on the test day over all other quadrants (Holm-Sidak, *p*<0.001). Within the vehicle group, the quadrant preference was not so striking, although they spent a higher amount of time in the reinforced quadrant as compared to the lateral quadrant A (*p*=0.009) and the contralateral D (*p=*0.033), but not to the other lateral quadrant B (*p*=0.38). When the amount of time swimming in the reinforced quadrant was compared between both groups, this result was confirmed, as the experimental rats spent more time searching in the previously reinforced quadrant than the vehicle rats (t=3.11, *p*=0.02) (Figure 3b).

#### 3.2. Metabolic activity increased in the dorsal DG and decreased in the ventral Ca1, IL cortex and mammillary bodies following spatial training under NPY Y_2_R blockade

Significant group differences in CCO activity were found in the prelimbic cortex (F_(2,21)_= 8.57; *p*=0.002), infralimbic cortex (F_(2,21)_=8.68; *p*=0.002), parietal cortex (F_(2,19)_= 8.68; *p*=0.002), CA1 region (F_(2,20)_= 4.67; *p*=0.023) and dentate gyrus (F_(2,20)_= 4.93; *p*=0.02) of the dorsal hippocampus. Additionally, statistical differences were found in the CA1 region of the ventral hippocampus (F_(2,21)_= 5.78; *p*=0.01), the central (F_(2,19)=_ 26.88; *p*<0.001) and lateral (F_(2,19)_= 3.81; *p*<0.05) nuclei of the amygdaloidal complex and the medial mammillary nucleus (F_(2,17)_= 12,45; *p*<0.001). Figure 4a, Figure 4b and Table 1 show the mean CCO activity values measured in the regions of interest of the different experimental groups, including a control cage (CC) group, used as a baseline for comparison.

**Figure 4:**
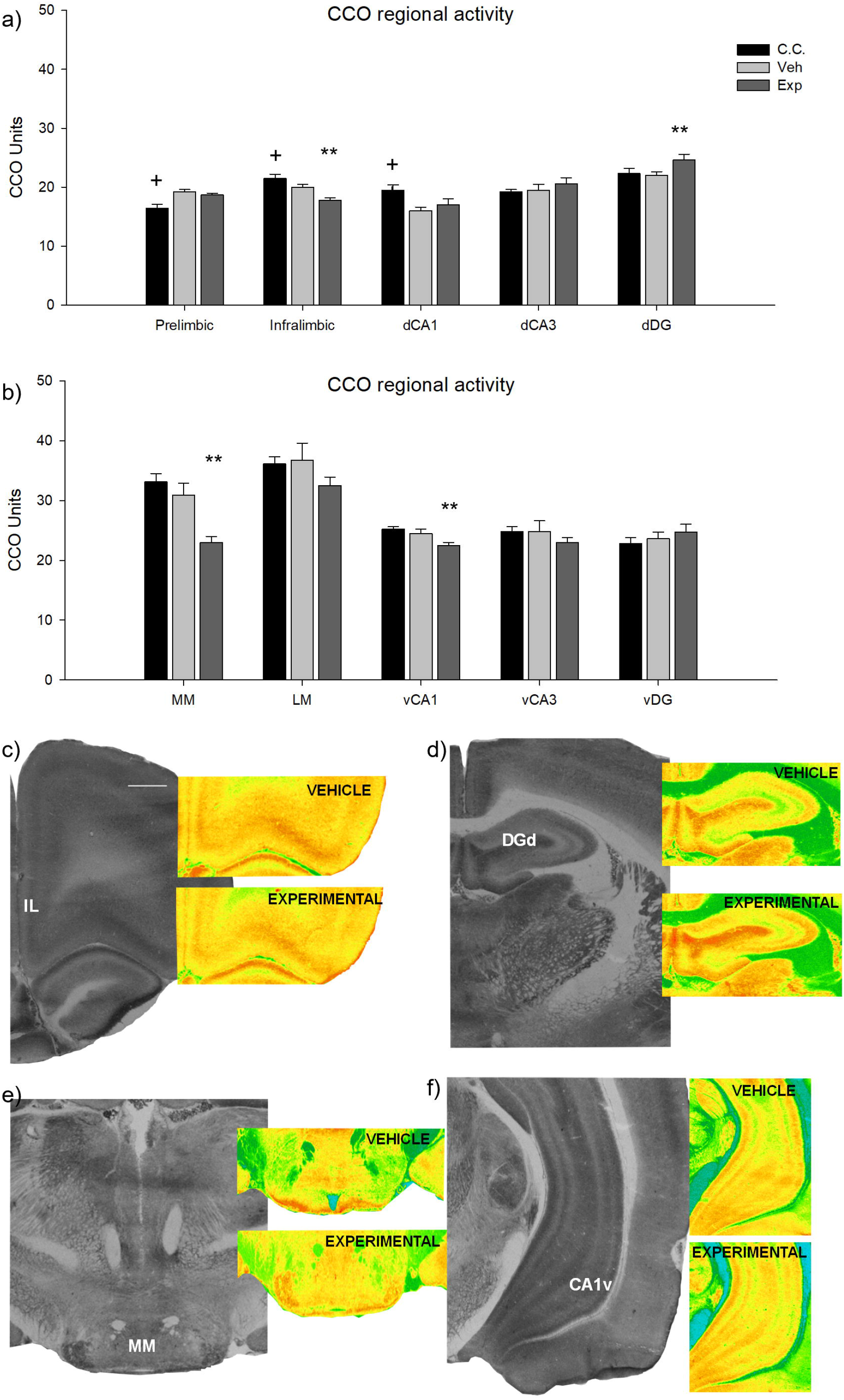
Brain metabolism was increased in the dorsal dentate gyrus and decreased in the ventral CA1, the mammillary bodies and the IL region of the mPFC after Y_2_R blockade prior to spatial memory retrieval. **a,b)** cytochrome c oxidase (CCO) activity results. Bars represent the mean ± S.E.M for each group. ** (*p*<0.05) experimental *vs* control and saline groups; ^+^ (*p*<0.01) control cage *vs* experimental group. **c-f)** Representative images of CCO histochemical stain performed in vehicle control and experimental groups showing: **c)** the prefrontal cortex (bregma 4.20 to 3.72 mm), **d)** dorsal hippocampus (bregma −2.64 mm to −3.72 mm), **e)** the mammillary bodies (bregma −4.2 mm to −4.68 mm) and **f)** the ventral hippocampus (bregma −4.8 mm to −5.28 mm) according to Paxinos and Watson (2004); for the exact values found in experimental, vehicle and control cage, please refer to Table 1. Black or red colours represent higher CCO activity, while green to blue colours symbolize lower activity. Scale bar: 1mm. The relative optical density of each region was measured by taking three non-overlapping readings in each section, in three consecutive sections by using a square-shaped sampling window adjusted for each region size (MCID, InterFocus Imaging Ltd., Linton, England). 150 x 150 mm (96 x 96 DPI). Abbreviations: CC (control cage group), Exp (Experimental group), Veh (Vehicle group), dorsal dentate gyrus (dDG), ventral (vDG), dorsal and ventral *Cornu Ammonis* 3 (dCA3, vCA3 respectively), dorsal and ventral *Cornu Ammonis* 1 (dCA1, vCA1 respectively), medial nucleus of the mammillary bodies (MM) and lateral nucleus (LM).

**Table 1.** Regional brain cytochrome c oxidase activity values. Mean ± S.E.M is shown for each experimental group. *(*p*<0.01) control cage vs experimental and Vehicle groups; ** (*p*<0.05) experimental *vs* control and saline groups; +(*p*<0.01) control cage *vs* experimental group and ++(*p*<0.05) control *vs* vehicle group.

|  | n | Vehicle | n | EXP | n | C. Cage |
| --- | --- | --- | --- | --- | --- | --- |
| **CORTEX** |  |  |  |  |  |  |
| Prelimbic | 8 | 19.2±0.4 | 6 | 18.7±0.3 | 8 | 16.4±0.7<sup>*</sup> |
| Infralimbic | 8 | 20.0±0.5 | 6 | **17.8±0.4**<sup>**</sup> | 8 | 21.5±0.7<sup>+</sup> |
| Cingulate | 8 | 19.8±0.7 | 6 | 19.2±0.5 | 8 | 21.2±0.9 |
| Motor | 8 | 20.6±0.4 | 6 | 19.9±0.3 | 8 | 20.5±0.4 |
| Parietal | 8 | 18.8±0.2 | 6 | 17.8±0.6 | 8 | 21.5±0.7<sup>+</sup> |
| **HIPPOCAMPUS** |  |  |  |  |  |  |
| CA1 dorsal | 8 | 16.0±0.6 | 6 | 17.0±1.0 | 7 | 19.5±0.9<sup>++</sup> |
| CA3 dorsal | 8 | 19.5±1.0 | 6 | 20.6±1.0 | 7 | 19.2±0.4 |
| DG dorsal | 8 | 22.0±0.6 | 6 | **24.6±1.0**<sup>**</sup> | 7 | 22.3±0.9 |
| CA1 ventral | 8 | 24.5±0.7 | 6 | **22.5±0.5**<sup>**</sup> | 8 | 25.2±0.4 |
| CA3 ventral | 7 | 24.8±1.8 | 6 | 23.0±0.8 | 7 | 24.8±0.8 |
| DG ventral | 8 | 23.6±1.1 | 6 | 24.7±1.3 | 7 | 22.8±1.0 |
| **DIENCEPHALON** |  |  |  |  |  |  |
| Medial Mammillary Nuclei | 6 | 30.9±2.0 | 6 | **23.0±1.0**<sup>**</sup> | 6 | 33.1±1.4 |
| Lateral Mammillary Nucleus | 6 | 36.7±2.9 | 6 | 32.5±1.4 | 6 | 36.1±1.2 |
| **AMYGDALOID COMPLEX** |  |  |  |  |  |  |
| Basolateral Nucleus | 7 | 21.7±0.8 | 6 | 21.1±1.6 | 7 | 20.6±0.9 |
| Medial Nucleus | 7 | 15.7±0.8 | 6 | 15.3±0.8 | 6 | 17.5±1.3 |
| Central Nucleus | 7 | 16.7±1.0 | 6 | 16.6±1.1 | 7 | 22.7±0.8<sup>*</sup> |
| Lateral Nucleus | 7 | 16.7±0.9 | 6 | 14.8±6.6 | 7 | 19.5±1.6<sup>+</sup> |

*EXP: Experimental group. C. Cage: Control Cage group. Vehicle: Vehicle group.*

In particular, Tukey’s HSD tests showed significant increases in CCO between groups trained in the water maze (Exp-Veh) as compared to the CC group; specifically, the activity of the prelimbic cortex was higher in Veh (*p*=0.021) and Exp (*p*=0.003) groups as compared to the CC group. In contrast, CCO activity was lower in the Veh and Exp groups in the central nucleus of the amygdaloid complex (*p*<0.001) as compared to the CC group. A similar effect was found in the lateral nucleus of the amygdaloid complex (*p*<0.001) in between the Exp and the CC group. Finally, the CA1 region of the dorsal hippocampus presented higher CCO activity in the CC group as compared to the Veh group (*p*=0.02).

On the other hand, the effect of the drug was observable in the differences found between Exp and Veh groups; specifically, significantly lower CCO activity was detected in the Exp group infralimbic region of the mPFC and the parietal cortex (*p*=0.002) (figures 4a and 4c). Additionally, significantly higher CCO activity (*p*=0.05) was found in the dentate gyrus of the dorsal hippocampus in the Exp group (figures 4a and 4d), whereas the ventral CA1 CCO activity was lower in the Exp than in the rest of the groups (figures 4b and 4f). Lastly, the mammillary nucleus of the Exp group showed significantly lower CCO activity (*p*<0.001) as compared to the rest of the groups (figures 4b and 4e).

## 4. DISCUSSION

The results of this study show that hippocampal NPY Y_2_R antagonism enhances spatial memory retrieval and alters retrieval-induced brain metabolic activity. Following retrieval under Y_2_R blockade, increased metabolic activity was detected in the DG of the dorsal hippocampus, and decreased activity was detected in the ventral CA1 hippocampal subfield, the medial mammillary bodies and the infralimbic region of the mPFC. Our findings support the notion that the NPY system is involved in spatial memory through the activation of Y2 receptors. Moreover, it suggests that Y_2_R-dependent changes in metabolic activity are a mechanism involved in spatial memory processing.

### 4.1. Intrahippocampal administration of an NPY Y_2_R antagonist rapidly improves the early recall of spatial reference memory

To evaluate the mnemonic effects of the NPY system in cognition, we used a hidden platform task in the MWM paradigm (Morris, 1984), the most used paradigm to study, in rodents, spatial memory, that is heavily dependent on intact hippocampal function (Méndez-Couz et al., 2015a; Méndez-Couz et al., 2015b; Zorzo et al., 2021a).

All groups of rats displayed similar decreasing latencies to reach the platform along training days and a preference for the previously reinforced virtual quadrant on the retention probe, showing that rats successfully acquired the initial spatial memory.

However, rats receiving intrahippocampal infusions of the Y_2_R antagonist performed better than those in the vehicle group, presenting lower latencies to reach the escape platform and more time spent in the reinforced quadrant during the retention probe. The results are consistent with differential sub-acute and acute effects of hippocampal NPY in the cognitive processes involved in the spatial reference memory task (Méndez-Couz et al., 2021). Likewise, other studies applying acute infusions of Y_2_R antagonists in the lateral ventricles (Kornhuber & Zoicas, 2017), dorsal hippocampus, amygdala or dorsolateral septum (Kornhuber & Zoicas, 2020) reported both Y1 and Y2 receptor-mediated NPY neurotransmission as necessary for retention and retrieval of hippocampus-dependent tasks, such as object recognition and retrieval of social memory. Together with our results, this would suggest the region-specific and memory type-specific role of NPY acting on Y1 or Y2 receptors on distinct phases of memory.

The NPY system is well known to be related to anxiety and mood regulation (Gøtzsche & Woldbye, 2016; Nwokafor et al., 2020; Thorsell et al., 2000; Vázquez-León, Ramirez-San Juan, Marichal-Cancino, Campos-Rodriguez, Chavez-Reyes, and Miranda-Paez et al., 2020). On the other hand, the role of Y_2_Rs in spatial versus emotional learning is still controversial (Hörmer et al., 2018; Tasan et al., 2016). In this regard, stress and locomotor activity could have affected the performance in the MWM test. In previous studies with this model (Méndez-Couz et al., 2021), we found, however, no anxiogenic or anxiolytic behavioural effects, ruling out the exclusively stress-related treatment influence on the subsequent MWM task. Additionally, previous studies in knockout Y_2_R (-/-) mice failed to show alterations in locomotor activity (Redrobe et al., 2004; Tschenett et al., 2003) or anxiety-related behaviour (Zambello et al., 2011). More recently, Hörmer et al. (2018) reported that the hippocampal Y_2_Rs modulate memory depending on its emotional valence, enhancing spatial memory while delaying fear conditioning memories. Taken together, these results suggest that the NPY Y_2_R blockade induced mnemonic-associated functional changes that are not exclusively attributable to anxiolytic or anxiogenic effects of the drug. Therefore, the observed regional metabolic changes (see below) are likely to be related to the specific effects of the treatment on retrieval of spatial memory.

### 4.2. Hippocampal NPY Y_2_R blockade before spatial memory retrieval changes metabolic activity in the dorso-ventral hippocampus, infralimbic cortex and mammillary bodies

The dynamic changes in brain metabolism in the dorsal and ventral hippocampus, mammillary bodies and prefrontal cortex after Y_2_R antagonist treatment were striking and may serve to explain the observed effects on behavioural performance.

We found a generalized reduction of CCO activity in the experimental group after retrieval. Decreased brain energy metabolic capacity has been recently related to a more efficient oxidative metabolism associated with underlying processes of synaptic plasticity, suggesting that a reduction in CCO activity is required for the stabilization of a preceding consolidated spatial cognitive mapping (Zorzo et al., 2021b). This might explain the better performance in the experimental group. However, the particular molecular mechanisms associated with the effects of Y_2_R blockade on regional brain oxidative metabolism are complex, involving the modulation of neuronal energy demands by the direct and indirect actions of NPY on different signalling systems, an issue that is difficult to disentangle. One possible mechanism is the action of NPY on other neurotransmitter systems (like catecholamines) and its regulation of glucose metabolism in both neurons and glial cells (Huang et al., 2021).

NPY Y_2_R has been implicated with several neuronal excitability (Silva et al., 2005) and cognitive processes (Botterill et al., 2015; Hörmer et al., 2018; Verma et al., 2019); this suggests a functional role on spatial orientation. Specifically, when the NPY is applied before the memory training, it seems to impair the acquisition and retention of spatial memory, apparently via presynaptic Y_2_R-mediated inhibition of synaptic glutamatergic transmission and LTP induction, [see review by Gøtzsche and Woldbye (2016)]. Y_2_R receptors are present among others, in the DG granular cells (Schwarzer et al., 1998), and strong evidence shows that this receptor undergoes experience-dependent metaplastic regulation (Méndez-Couz et al., 2021), shifting from high affinity to low-affinity states of this receptor (Parker et al., 2007). The consequence of the inhibition of Y_2_R receptors at glutamate terminals in the hippocampus would therefore be the rise in the glutamate release, but it remains unclear how this could support the improved hippocampus-dependent spatial learning. Under physiological conditions, the suppression of glutamate release by NPY acting on Y_2_R receptors goes against information encoding through long-term potentiation. In turn, this blockade would favour the long-term depression (LTD), an intrinsic component of the acquisition of complex representations (Kemp & Manahan-Vaughan, 2004, 2007, 2008), which has been related to improvement in the water maze performance (Dong et al., 2013). The Y_2_R-mediated LTD would be associated with higher synaptic plasticity and the observed retrieval improvement in the water maze task after the Y_2_R blockade. In agreement, spatial memory in the Barnes test was enhanced after Y_2_R suppression; moreover, after hippocampal reactivation in a Y_2_R knockout rat model, the improved spatial memory was reduced (Hörmer et al., 2018).

Remembering a past event involves the reactivation and partial modification of the patterns of neural activity present at encoding (Richards & Frankland, 2017). In this regard, we had previously reported a modified mPFC-dorsal hippocampus functional coupling involvement during the acquisition and retrieval of a spatial memory task (Conejo et al., 2010; Conejo et al., 2013; Méndez-Couz et al., 2015a).

We found that blockade of Y_2_R receptors prior to a spatial memory retrieval test increases the dDG metabolic activity. The dDG receives projections from the medial entorhinal cortex (Wyss, 1981) and forms part of the circuits encoding for the “what” and “where” aspects of context-dependent learning (Hoang et al., 2018). Recently, it has been proposed that the DG is functionally involved in the discrimination between temporal and spatial experiences, due to the strong segregation of information encoding in its upper and lower blades (Hoang et al., 2018; Strauch & Manahan-Vaughan, 2020). In a contextual-dependent spatial task performed in the T-maze (Méndez-Couz et al., 2019), increased IEG expression in the dDG lower blade was reported, consistent with its recruitment by the recall of context-dependent or “where” information encoding. Converging evidence suggests that dentate gyrus cell interconnectivity is also essential for memory recall and its precision over time (Haubrich & Nader, 2018). Taken together, the observed metabolic state change in the DG of experimental rats is consistent with the DG critical involvement needed for an effective and precise recall of an already known place (Emerich & Walsh, 1989; Méndez-Couz et al., 2015a; Zorzo et al., 2021a).

A significant decrease of the ventral CA1 metabolic activity was found after dorsal hippocampal Y_2_R antagonism. Whereas the dorsal portion of the hippocampus is mainly related to cognitive processing, the ventral pole has been traditionally associated with emotional and bodily states (Fanselow & Dong, 2010). Therefore, one could be tempted to interpret the differences in CCO activation found in this area as those typically related to the drug-infusion effect on anxiety or stress responses. However, consistently with the lack of anxiogenic or anxiolytic effects in this model (Méndez-Couz et al., 2021), no differences in mean CCO activity were found in brain regions typically associated with stress and anxiety, like the amygdala (Villarreal et al., 2002). Although the role of the ventral CA1 in spatial memory is not entirely clear, these metabolic results would agree with previous metabolic, electrophysiological and IEG studies pointing towards the involvement of the ventral hippocampus in rewarded spatial memory (Beer et al., 2014; Méndez-Couz et al., 2016; Sosa et al., 2020).

Besides, our results showed a metabolic activity decrease in the mammillary bodies after hippocampal Y_2_R blockade. This diencephalic structure is crucial for transmitting information to higher forebrain centres via the mammillothalamocortical pathway, being involved in the neural circuitry for the recollection of memories (Guillery, 1955; Vann, 2005, 2010). Consistently, lower activity in the MM was also found after spatial memory extinction of a task previously acquired in the MWM (Méndez-Couz et al., 2014; Méndez-Couz et al., 2016), which is not surprising, taking into account that the mammillary bodies have been conventionally considered a hippocampal relay. Indeed, spatial memory deficits were observed after lesions in the mammillary bodies or their major efferent, the mammillothalamic tract (Vann, 2010).

Lastly, a lower CCO activity was observed in the infralimbic region of the mPFC. We have previously found decreased levels of Y_2_R and increased Y_1_R expression in the mPFC, following hippocampal Y_2_R antagonism and after the MWM task (Méndez-Couz et al., 2021). These results were not surprising, given that the continued and intrinsic increase in hippocampal excitability changes the excitation-inhibition equilibrium in the mPFC by modifying GABA receptor expression (Grüter et al., 2015). Additionally, the Y_2_R antagonism causes disinhibition of glutamatergic and noradrenergic terminals in the hippocampus that might change the excitatory output from the hippocampus to the mPFC. This, in turn, may have triggered the changes in the metabolic activity of the IL cortex that we detected in the experimental group.

At a metabolic level, changes in the mPFC have been found at the late stages of acquisition (Conejo et al., 2010), retrieval (Zorzo et al., 2021a; 2021b) and the extinction of spatial memory tasks in the water maze (Méndez-Couz et al., 2014, 2015b). During early recall, the metabolic activation in the dDG of the hippocampus is negatively correlated with the metabolic levels measured in the IL cortex (Méndez-Couz et al., 2015a), which would fit with our results in the experimental group, showing a better performance associated with a higher DG and a lower IL metabolic energy levels. However, given the mediation of the mPFC in the attentional and motivational components of the spatial orientation in the MWM (Conejo et al., 2007; Conejo et al., 2010), one cannot rule out the possibility of motivational aspects influencing a better retrieval. In line with this interpretation, the IL cortex is reported to modulate appetitive Pavlovian extinction (Mendoza et al., 2015) and extinction learning of a spatial memory task (Méndez-Couz et al., 2014), in which the persistence of the previously reinforced learned response needed to be decreased. This aspect of mPFC function, together with the above-mentioned rapid NPY metaplastic changes in the hippocampal-mPFC circuit, may account for the improved spatial memory recall observed in experimental rats after the Y_2_R antagonist administration.

Our results help to further illuminate the role of the NPY system in spatial learning. We demonstrate that under hippocampal Y_2_R antagonism, spatial memory recall is enhanced and that such enhancement is correlated with changes in regional brain energy metabolism along the dorso-ventral axis of the hippocampus, the medial mammillary bodies and the infralimbic region of the mPFC. Taken together, these results suggest that Y_2_R exert control over patterns of brain activation that are relevant for spatial memory expression.

## Author Contributions

The study was designed by NC and MM-C. Experiments were conducted by MM-C, HGP and NC, and analysed by all authors. MM-C and NC interpreted the results and wrote the article.

## Acknowledgements

We gratefully thank Prof. Ana Paula Silva for providing the Y_2_R receptor antagonist.

## Funding Sources

This work was supported by the following grants: MINECO, Spain [grant numbers PSI2017-83038-P, PSI 2017-83893-R and PSI2017□90806□REDT].

## Statement of Ethics

All experimental procedures carried out with animals were approved by the local Animal Ethics Committee of the University of Oviedo and following the European Communities Council Directive 2010/63/UE and the Spanish legislation on care and use of animals for experimentation (Royal Decree 53/2013). All efforts were made to minimize the number of animals used and their suffering.

## Notes

### Competing Interest Statement

The authors have declared no competing interest.

